# Effects of blue monochromatic light directed at one eye of pregnant horse mares on gestation, parturition and foal maturity

**DOI:** 10.1101/2021.07.09.451773

**Authors:** Anne Lutzer, Christina Nagel, Barbara A. Murphy, Jörg Aurich, Manuela Wulf, Camille Gautier, Christine Aurich

## Abstract

Blue light directed at one eye advances the equine ovulatory season but can also advance foaling. In this study, effects of blue LED light on pregnancy outcome were assessed. Twenty mares with singleton pregnancies were studied over two consecutive years in a cross-over design. In one year, mares received an extended photoperiod using 50 lux of blue LED light (468 nm) directed at a single eye from 08:00 until 23:00 daily via head-worn light masks starting mid-December and in the other year remained untreated as controls. Gestation was shorter in blue LED light-treated (333.2±1.0 days,±SEM) than in control pregnancies (337.9±1.8 days, p<0.05). Colostral IgG content was lower in treated than in control pregnancies (p<0.05) but was adequate to ensure sufficient IgG transfer to newborn foals with a single exception. Foals born to blue LED light-treated mares had lower wither heights (102.5±1.1 vs. 105.2±1.0 cm, p<0.01), similar weights (55.0±1.3 vs. 55.5±1.1 kg) and took less time to stand after birth than control foals (39±3 vs. 56±4 min, p<0.05). The neutrophil/lymphocyte ratio, was higher in foals born to blue LED light-treated mares than controls (3.2±0.2 vs. 2.7±0.2; p<0.05). Foals born to blue LED light-treated mares had reduced hair length compared to controls (13.1±0.8 vs. 20.9±0.1 mm; p<0.001) and hair regrowth in treated mares was reduced (p<0.05). Maternal plasma prolactin concentration was increased in treated mares (p=0.05) and peaked at foaling (p<0.001). In conclusion, blue LED light directed at one eye advanced foaling and influenced height, hair coat and maturity in foals.

## Introduction

The horse is a seasonally breeding species and extending day length by exposing mares to artificial light for several hours after dusk from early December onwards advances the onset of seasonal reproductive activity. This is achieved via shortening the daily phase of melatonin release from dusk to dawn (reviewed by Palmer and Guillaume, 1992). Short wavelength light within the blue spectrum is particularly effective at suppressing melatonin secretion in the horse and low intensity blue light from light emitting diodes (LED) directed at a single eye has been used to advance the ovulatory season in mares to the same extent as illuminating the mares’ stable (Walsh et al., 2013; Murphy et al., 2014).

Photoperiod also affects equine pregnancy. Gestation length has been reported to decrease as the daily light phase increases throughout the foaling season (Hintz et al., 1979). In contrast to a longer gestation, however, foals born early in the year were smaller than foals born later in the foaling season but did not differ in weight (Beythien et al., 2017). In agreement with a physiologically shorter gestation when day length increases in spring, artificial light programmes (Hodge et al., 1982) and blue LED light directed at one eye of pregnant mares can shorten gestation length (Nolan et al., 2017) compared to control mares maintained under ambient natural photoperiods.

It has been suggested that light programmes applied to pregnant mares may stimulate final fetal maturation and growth (Nolan et al., 2017) but results so far remain ambiguous. In a study using conventional light programmes in pregnant mares, a numerical weight difference between foals born to light-exposed and control mares was reported, however, statistical significance was nor reached (Hodge et al., 1982). In a more recent project, the effects of blue LED light directed at one eye of the mare were studied. A decreased gestation length but similar foal birth weight was demonstrated in one of the reported experiments where animals were selected based on a previous history of prolonged gestation lengths (>350 days). A second experiment used mares bred to one of two stallions and gestation length was not affected by administration of pre-partum extended blue LED light, whereas foals born to treated mares had higher birth weights (Nolan et al., 2017). Foal size was not determined in these studies but may be considered a more important predictor of performance potential than birth weight for some equestrian disciplines. The precise effects of blue LED light on equine gestation and fetal growth thus need to be verified.

Interestingly, blue LED light treatment of pregnant mares was associated with a shorter hair coat in the foals born to treated mares compared to foals from control mares (Nolan et al., 2017). The onset of vernal hair shedding in mares is mediated by the anterior pituitary hormone prolactin (Thompson et al., 1997). Prolactin concentration in plasma of mares follows an annual rhythm with a peak in summer (Johnson, 1986). Hair shedding has been induced by experimental treatment with prolactin (Thompson et al., 1997). Treatment of mares with melatonin in spring, simulating continuation of a short daily light phase, reduced plasma prolactin concentration and delayed hair shedding (Aurich et al., 1997). Furthermore, in pregnant ewes, artificial long daylight programmes providing a summer photoperiod or constant light increased plasma prolactin concentrations in ewes and in their newborn lambs (Bassett, 1992; Parraguez et al., 1996). Effects of blue LED light on fetal hair growth (Nolan et al., 2017) might therefore be mediated by changes in maternal prolactin release.

This study aimed to evaluate the effects of pre-partum administration of extended daily blue LED light on equine pregnancy, foaling, foal maturity and viability. We hypothesized that blue LED light applied for 15 hours per day would shorten gestation length, stimulate foal development and reduce hair growth via an increase in maternal plasma prolactin concentration. Because 90% of horse mares give birth at night and only 10% foal during daylight hours (Rossdale and Short, 1967; Heidler et al., 2004), we also investigated if blue LED light treatment shifted the timing of foaling.

## Materials and Methods

### Animals

The study was approved by the competent authority for animal experimentation in Brandenburg State, Germany (*Landesamt für Arbeitsschutz, Verbraucherschutz und Gesundheit*, license number 2347-41-2018).

A total of 20 Warmblood sport horse (*Equus caballus*) mares with singleton pregnancies were available for the study over two consecutive breeding seasons. All animals were broodmares of the Brandenburg State Stud at Neustadt (Dosse), Germany (52° 52′ 0″ north, 12° 25′59″ east, at winter solstice sunrise 8:20, sunset 15:56). At the beginning of the study, mares were 8.2±1.1 years old (range 3-17 years). Mares were bred by artificial insemination to different Warmblood stallions selected by the stud management. For management reasons, breeding of mares was limited to the time between March and June resulting in a short foaling season in the following year. Day of pregnancy was calculated from the date of ovulation, which had been checked at daily intervals by transrectal ultrasonography before and after insemination of the mares. During the summer months, i.e. in early and mid-pregnancy, mares were kept on pasture with access to a barn during nights or periods of extreme heat. Depending on the vegetation, mares were fed hay in addition to pasture grass. During winter and spring, i.e. the experimental period of our study, all mares irrespective of treatment were housed in the same group stable on straw and had daily access to an outdoor paddock. They were fed oats and hay twice daily. For feeding of oats, mares were tethered at their individual feeding places. Mineral supplements and water was freely available at all times. Approximately 14 days before the calculated date of parturition, mares were transferred to single boxes in the foaling unit. From this time on they were observed near-constantly until foaling. In the foals, time from birth to first standing and to first suckling their dam’s udder was recorded. The exact time to first standing was not available for one foal from the blue LED light group and exact time to first suckling was not available for one foal from each group.

### Experimental design

Mares were available for two consecutive years and only mares pregnant in both years were included into the study. In one year, mares received an artificially extended photoperiod using 50 lux blue LED light (468 nm) directed at a single eye from 8:00 until 23:00 daily via individual head worn light masks (Equilume, Kildare, Ireland) as described previously (Nolan et al., 2017) with some modifications. The Cashel mask model used in the present study delivers light for 15 h per day ensuring a constant delivery of blue-enriched light throughout the lengthened day while the model used by Nolan et al. (2017) delivered blue light only from 16:00 to 23:00 daily. In the other year, mares served as their own controls and received no light masks and thus were exposed to light only between natural sunrise and sunset. Light treatments were started on 13 December in year 1 and 16 December in year 2, corresponding to day 234±6 of pregnancy. At winter solstice, the time between sunrise and sunset was 7 h 43 min. On the days of the first (24 February) and the last foaling in control pregnancies (4 May), the time between sunrise and sunset was 10 h 35 min and 15 h 6 min, respectively.

Initially, 31 pregnant mares had been available in the first year and were ranked by expected foaling date and age and allocated in alternating order to blue LED light (n=15) and control treatments (n=16). Out of the 31 mares, in the second year 11 were no longer available because they did not become pregnant, were retired from breeding or sold, leaving 20 mares to be included into analysis for the study. Out of these 20 mares, eight mares received light masks in year 1 and served as controls in year 2 while 12 mares received light masks in year 2 and control treatments in year 1.

**Table 1:** Reproductive history of mares when entering the study and sex ratio of foals in blue LED light and control pregnancies (n=20)

|  | Light | Control |
| --- | --- | --- |
| Age (years) | 8.8 ± 1.1 | 8.6 ± 1.1 |
| Number of previous foals (n) | 4.8 ± 0.9 | 4.6 ± 0.9 |
| Sex ratio of foals (male/female) | 7/13 | 9/11 |

*No significant differences between light and control pregnancies*

### Mare weight, foal size and weight and immunoglobulin G in colostrum and foal plasma

Weight of mares was determined at regular intervals with an electronic horse scale (PW Bosche, Damme, Germany). The weight on days 284, 299 and 315 of pregnancy, day 3 before and days 1 and 6 after foaling was compared. At foaling, immunoglobulin G (IgG) content in colostrum was determined by sugar refractometer as described previously (Chavatte-Palmer et al., 2001). Colostrum was collected from both mammary complexes of each mare and the mean of both measurements used for further comparisons. Placental weight was recorded directly after shedding of the placenta and the placental surface area was measured as described (Allen et al., 2002; Beythien et al., 2017).

The IgG content in foal plasma was measured by commercial semiquantitative on-site test (Snap Foal IgG, Idexx, Kornwestheim, Germany) at approximately 18 hours after birth. This test classifies IgG content as insufficient (<400 ng/ml), intermediate (400-800 ng/ml) and adequate (>800 mg/dl). One day after birth, foals were weighed and the following body size parameters were determined as described previously (Beythien et al., 2017): height at withers, chest circumference, distance fetlock to carpal joint, distance carpal joint to elbow and cannon bone circumference.

### Progestagen and prolactin analysis

Blood for analysis of progestagens and prolactin in plasma was collected from one jugular vein into heparinised tubes (Vacuette, Greiner, Kremsmünster, Austria). Progestagen and prolactin concentration was determined on days 11 and 4 before foaling, within 30 min after foaling and on days 1 and 6 after foaling. Blood samples were centrifuged (1200 g, 10 min) and plasma was stored at −20°C until analysis.

Progestin concentration was analysed by enzyme-linked immunosorbent assay (Enzo Progesterone ELISA, Enzo Life Sciences, Farmingdale, NY, USA) as described previously (Beyer et al. 2019). The intra-assay coefficient of variation was 5.6%, the inter-assay coefficient of variation was 8.0% and the minimal detectable concentration 9.5 pg/ml.

Prolactin concentration was determined with an enzyme-linked immunosorbent assay developed for prolactin analysis in human plasma (DE1291, Demeditec Diagnostics, Kiel, Germany) following the protocol provided by the manufacturer. Serial dilution of equine plasma resulted in changes in prolactin concentration parallel to the standard curve and recovery of prolactin standard added to equine plasma was 90%. The intra-assay coefficient of variation was 15.9%, the inter-assay coefficient of variation was 7.9% and the minimal detectable concentration calculated as two standard deviations from zero binding was 420 pg/ml.

### Hair coat

Length of the guard hair and hair re-growth in mares were measured on day 1 after foaling as described previously (Schrammel et al., 2016). For assessment of re-growth, an 8 by 6 cm rectangular area was shaved on the left side of the chest at the beginning of the study in December and length of subsequently growing hair was measured. Length of the undisturbed guard hair was determined at the shoulder in the middle third of the spina scapula, using 0.1 cm increments. Hair including the root was torn out, placed on a sheet of paper and length of at least five single hairs determined with a flexible ruler and the mean length calculated. Hair length of neonatal foals was determined as described for the mares.

### Hematology

In the foals, blood for haematology was collected into EDTA-containing tubes (Vacuette, Greiner) within 30 min after birth. A complete blood count was performed using routine techniques as described previously (Melchert et al., 2019). As a measure of foal maturity (Jeffcott et al., 1982) the ratio of neutrophil granulocytes to lymphocytes (N/L ratio) was calculated. Besides the N/L ratio, data for total leukocyte count are presented.

### Statistical analysis

Statistical comparisons were made with the IBM-SPSS 26 statistics package (IBM, Armonk, NY, USA). Parameters determined repeatedly (weight, progestagen and prolactin concentration in mares) were compared by analysis of variance using a general linear model (GLM) for repeated measures with both time and treatment as within subject factors, taking into account that data from the same mares were compared for two consecutive years. Parameters determined only at one time were compared between treatments (blue LED light and control) by Wilcoxon test. Time of foaling throughout the day was grouped into four intervals of six hours each and frequency distribution compared between blue LED light treatment and control pregnancies by χ^2^-test. For all comparisons, a p-value <0.05 was considered significant. All data given are means ± standard error of mean (SEM).

## Results

All mares and their foals were healthy throughout the study, and no complications of pregnancy or parturition occurred. Body weight of mares increased with advancing gestation and decreased as expected post foaling (p<0.001) but at no time differed significantly between blue LED light and control pregnancies (figure 1).

**Figure 1:**
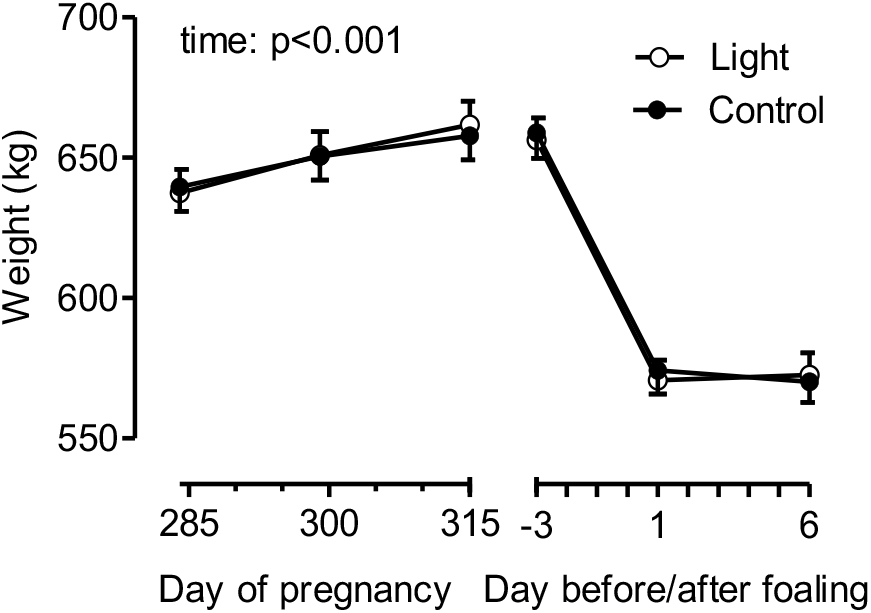
Body weight of mares (n=20) receiving blue LED light and control treatments from day 284 of pregnancy to day 6 after foaling, results of statistical analysis are indicated in the figure.

Gestation length differed between blue LED light and control pregnancies and was shorter when mares were equipped with light masks (light 333.2±1.0, control 337.9±1.7 days, p<0.05, figure 2a). Time from initiation of blue LED light treatment to foaling was 98.8±5.9 days. The IgG content indicated by the sugar refractometer BRIX was lower in colostrum of mares equipped with light masks compared to control pregnancies (p<0.05, figure 2b). Neither placental size nor placental surface differed between blue LED light pregnancies and control pregnancies (figure 2c,d).

**Figure 2:**
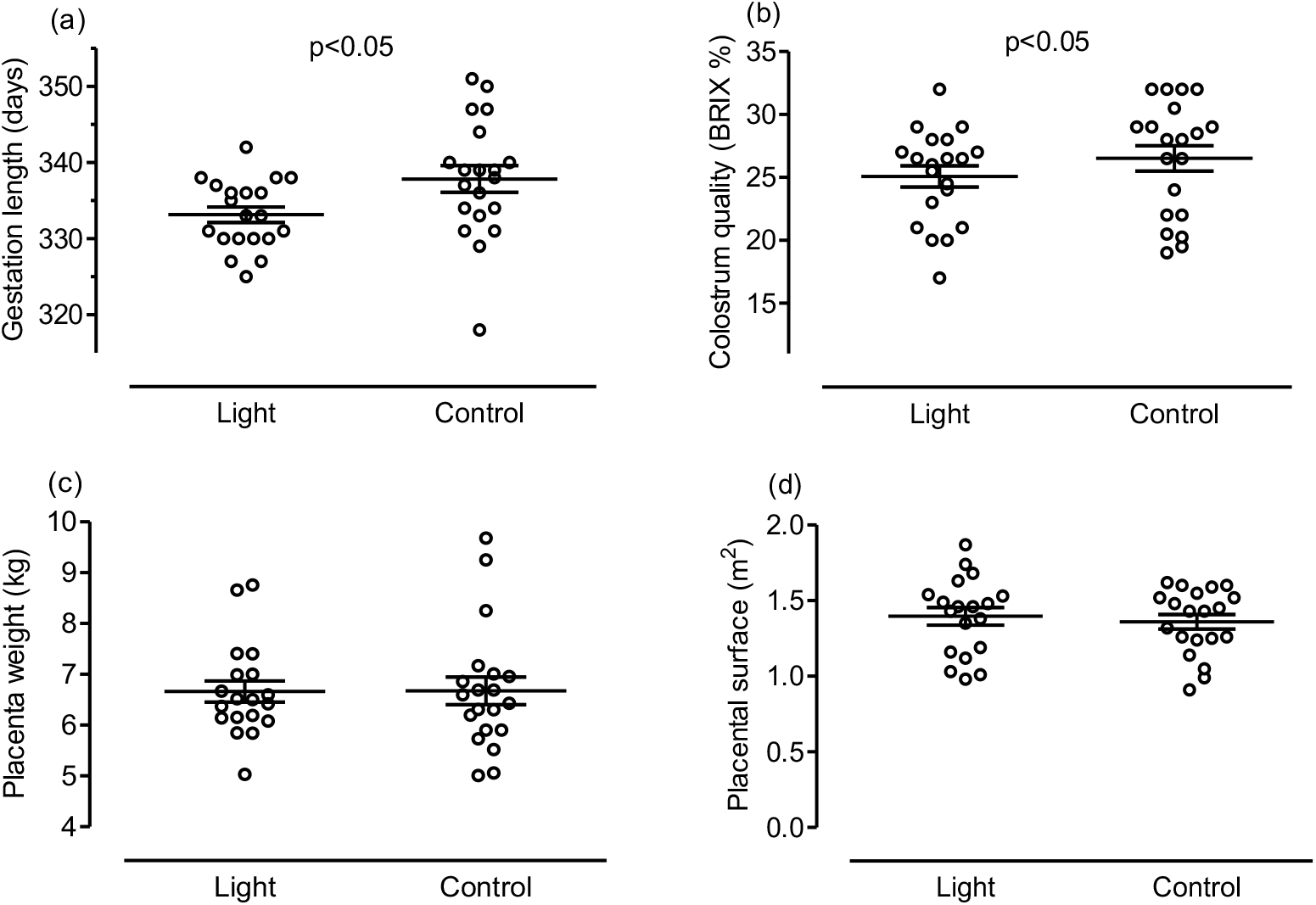
(a) Gestation length, (b) colostrum quality (BRIX), (c) placental weight and (d) placental surface in mares (n=20) receiving blue LED light and control treatments, results of statistical analysis are indicated in the figure.

Foaling occurred more often during the night (18:00 - 6:00) than during daytime (6:00-18:00) but the frequency distribution did not differ significantly between pregnancies when mares received blue LED light and control pregnancies (figure 3).

**Figure 3:**
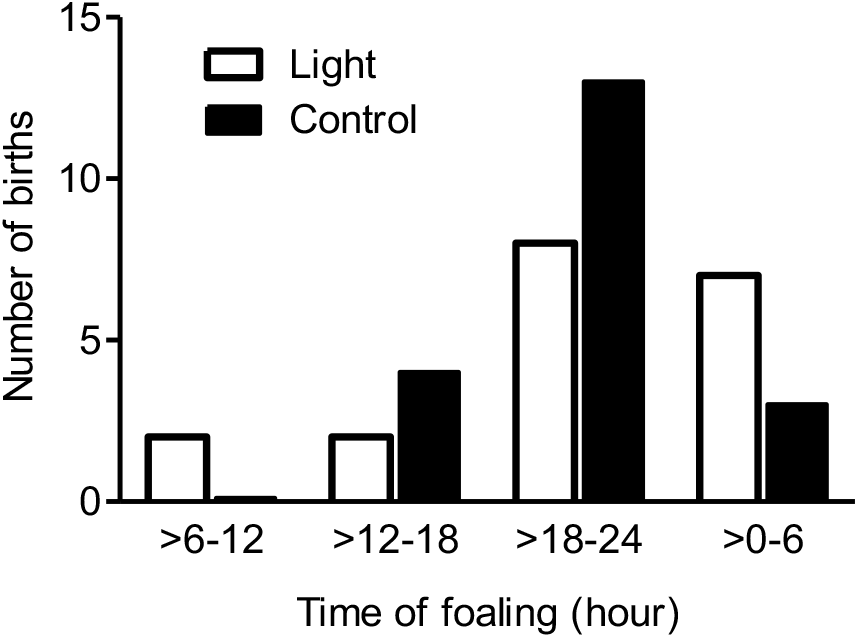
Time of foaling during the day in mares (n=20) receiving blue LED light and control treatments, no significant differences between treatment and control pregnancies.

On day 1 after birth, foals born to mares exposed to blue LED light had lower height at withers (light 102.5±1.1, control 105.2±1.0 cm p<0.01, figure 4a) and reduced elbow to carpus distance (light 30.6±0.3, light 31.8±0.3 cm, p<0.05, figure 4b) than control foals while there was no group difference in the carpus to fetlock distance (light 27.6±0.2, control 27.7±0.2 cm, figure 4c). Cannon bone circumference tended to be smaller in foals born to blue LED light-treated mares than in control foals (light 13.4±0.2, control 13.8±0.2 cm, p=0.061, figure 4d). Body weight (light 55.0±1.3, control 55.5±1.1 kg) and chest circumference (light 83.7±0.6, control 84.1±0.6 cm) were close to identical in foals born to mares exposed to blue LED light and foals from control pregnancies (figure 4e,f).

**Figure 4:**
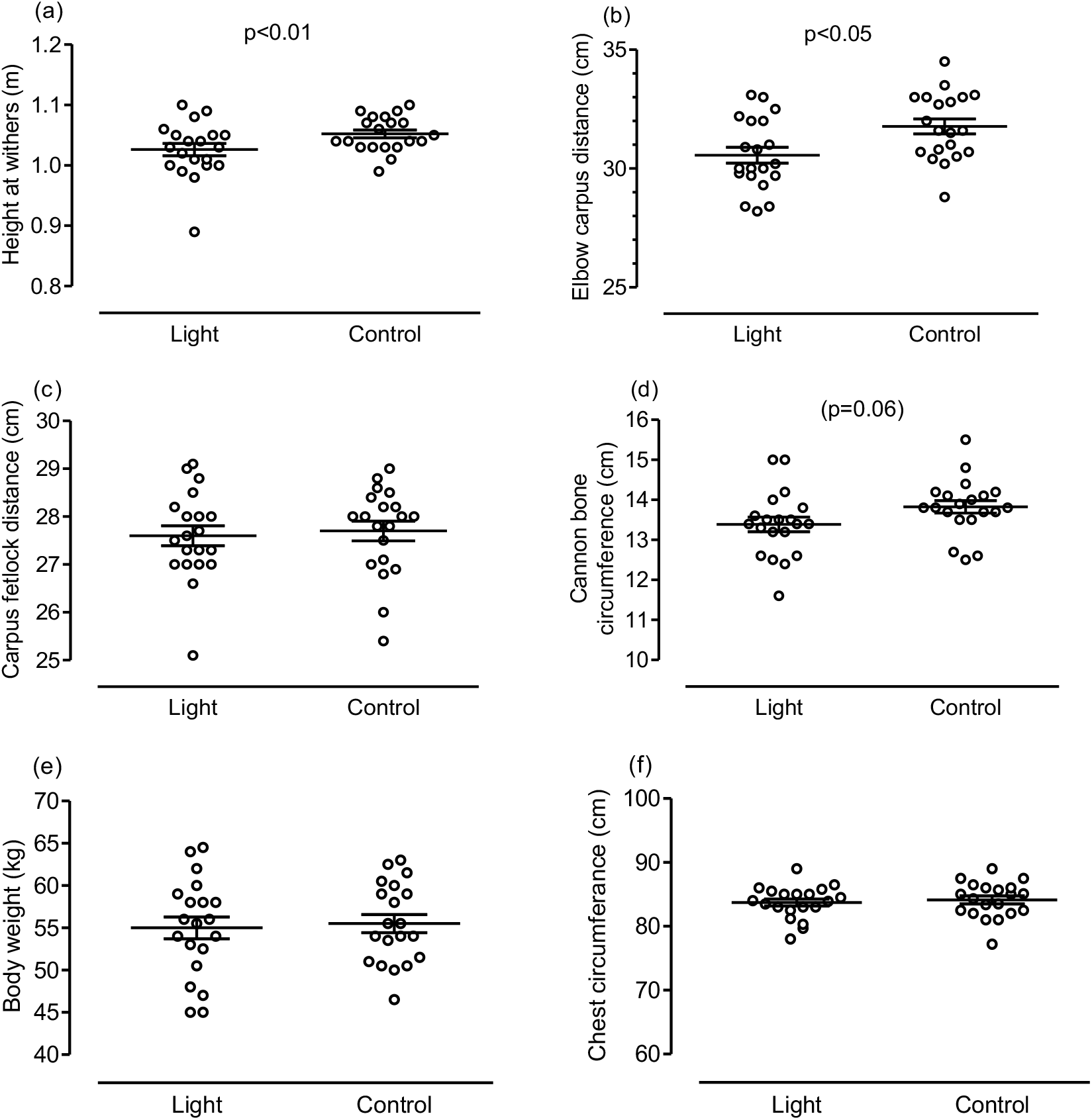
(a) Height at withers, (b) elbow to carpus distance, (c) carpus to fetlock distance, (d) circumference of the cannon bone, (e) body weight and (f) chest circumference in foals (n=20) born to mares receiving blue LED light and control foals (n=20), results of statistical analysis are indicated in the figure.

Progestagen concentration decreased during the last four days before foaling and further until day 6 after foaling (p<0.001) but changes in plasma progestagen concentration were close to identical when mares received blue LED light or were untreated (figure 5a). Prolactin concentration in plasma of mares increased steadily over 10 days before parturition, peaked on day 1 after foaling and decreased rapidly thereafter (p<0.001). For the time period from day -11 to the day of foaling (day 0), prolactin concentration in maternal plasma tended to be higher in blue LED light compared to control pregnancies and statistical significance was nearly reached (p=0.05, figure 5b).

**Figure 5:**
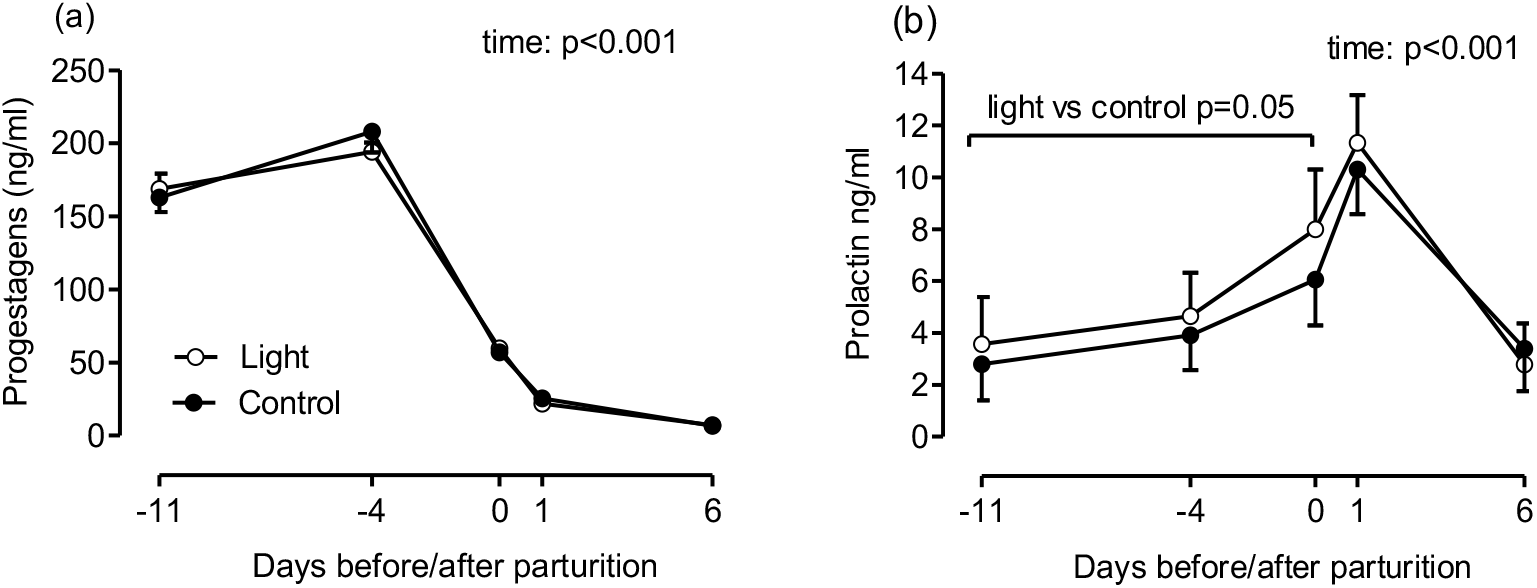
(a) Progestagen and (b) prolactin concentration in plasma of mares (n=20) receiving blue LED light and control pregnancies, results of statistical analysis are indicated in the figure.

Hair length in mares was not significantly changed by blue LED light treatment (figure 6a) but hair re-growth in a previously shaved area was less when mares were exposed to blue LED light (light 5.2±0.2, control 6.9±0.5 mm, p<0.05, figure 6b). Hair length of newborn foals was clearly reduced when mares had been exposed to blue LED light during gestation compared to foals from control pregnancies (light 13.1±0.8, control 22.9±0.9 mm, p<0.001, figure 6c).

**Figure 6:**
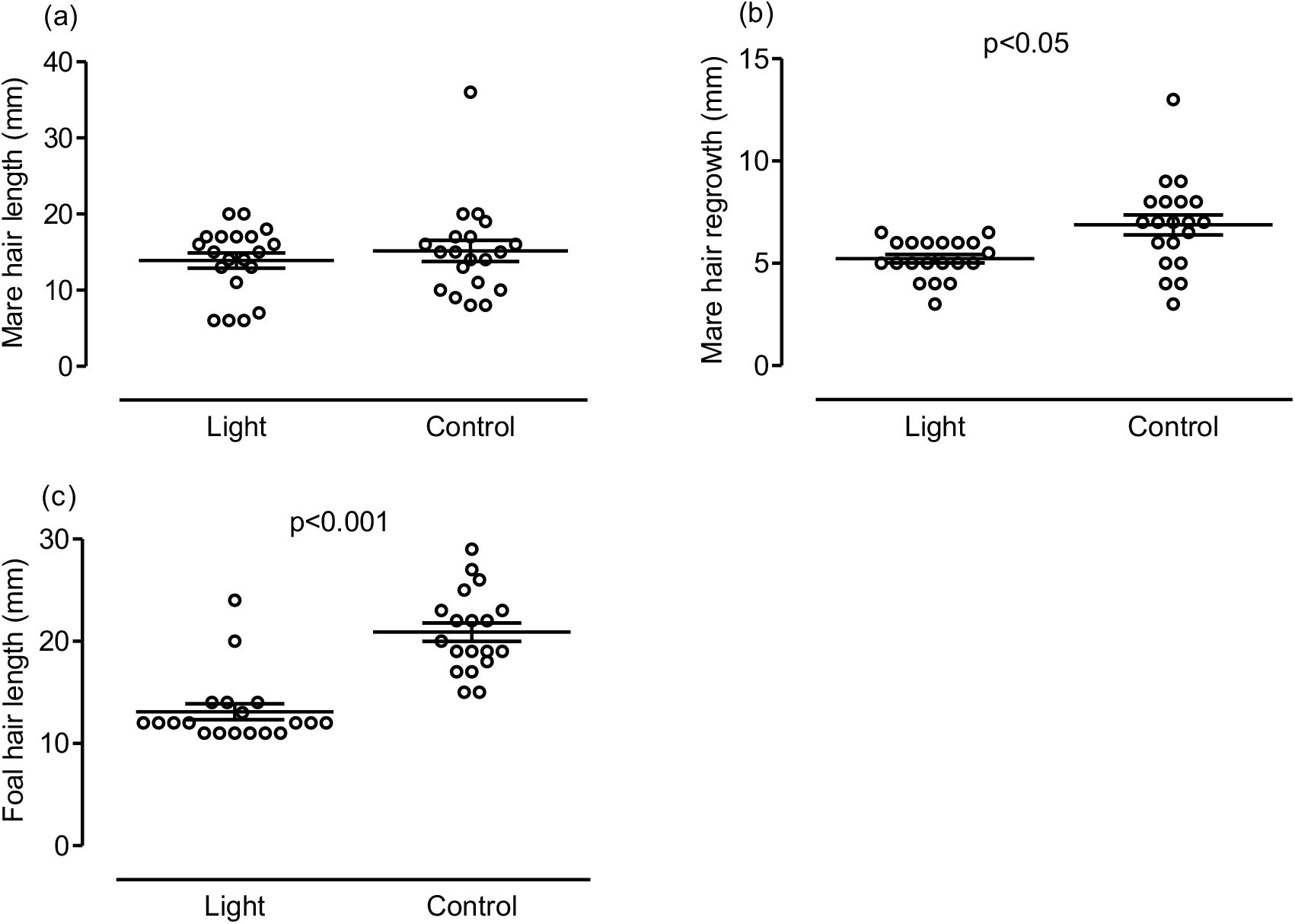
(a) Hair length and (b) re-growth of shaved hair in mares (n=20) receiving blue LED light and control treatments (c) hair length in foals born to mares receiving blue LED light and control foals, results of statistical analysis are indicated in the figure.

Foals born to mares exposed to blue LED light stood earlier after birth than control foals (light 39±3, control 56±4 min, p<0.05) while there was no difference between foal groups in time from birth to first suckling their dams’ udders (figure 7a,b). Total leukocyte count at birth did not differ between foals born from blue LED light-treated pregnancies and control foals (figure 7c), but the N/L ratio in foals directly after birth was higher when their dams had been exposed to blue LED light (light 3.2±0.2, control 2.7±0.2, p<0.05, figure 7d). All except one foal had an IgG concentration >800 ng/ml. In this foal from the blue LED light group, IgG concentration in plasma 18 h after birth was below 400 ng/ml.

**Figure 7:**
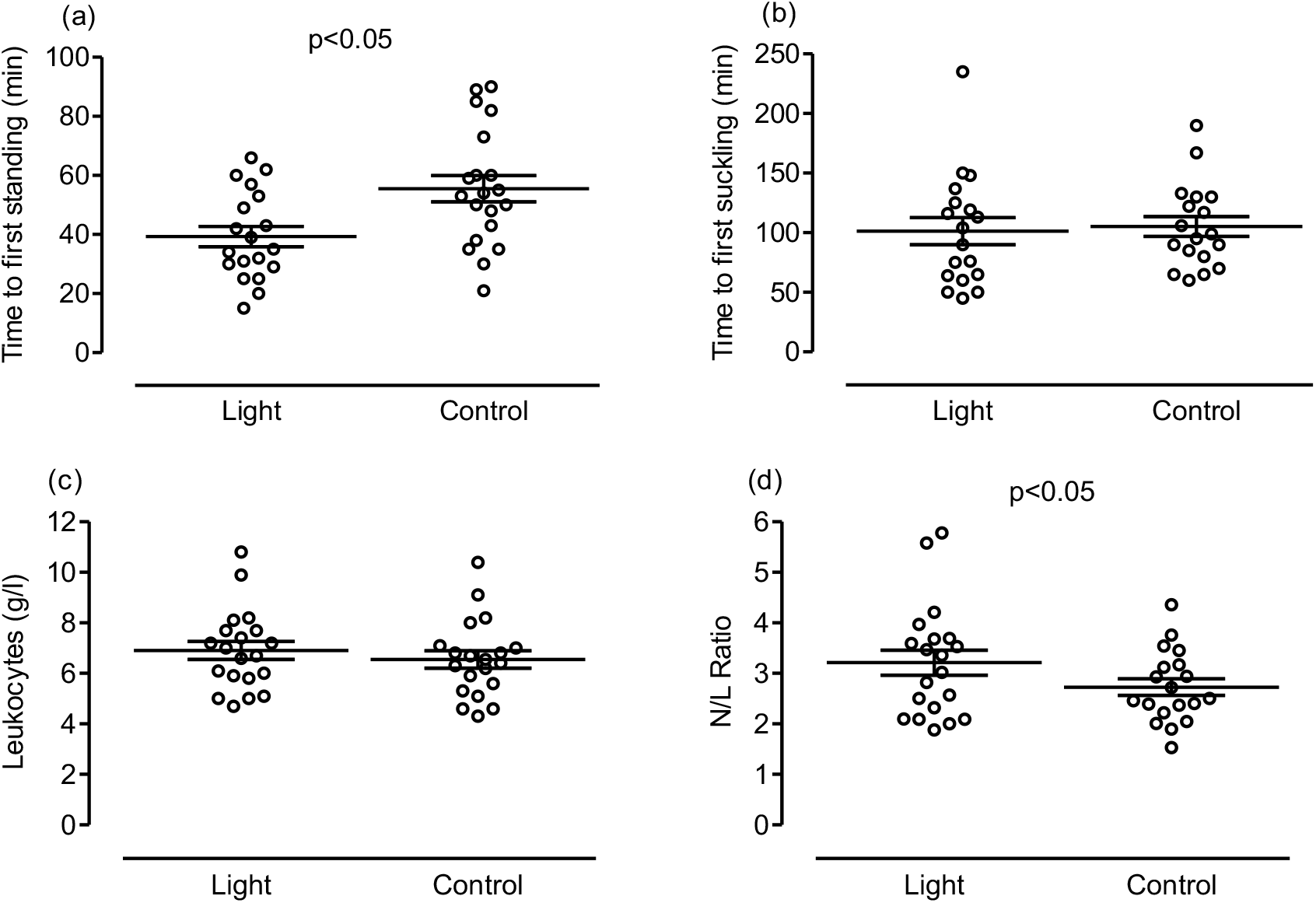
(a) Time from birth to first standing, (b) time from birth to first suckling, (c) leukocyte count and (d) neutrophil to lymphocyte (N/L) ratio in foals born to mares receiving blue LED light (n=20) and control foals (n=20), results of statistical analysis are indicated in the figure.

## Discussion

Pre-partum extension of day length using blue LED light had clear effects on pregnancy in mares and their newborn foals. Mares exposed to a long day photoperiod administered by way of a light mask delivering blue LED light to one eye from mid-December onwards foaled earlier than the same mares in their control pregnancies. Foals born to mares exposed to blue LED light were smaller at birth and had a markedly shorter hair coat but did not differ in weight from foals born from control pregnancies.

An effect of blue LED light applied from sunset until 23:00 on gestation length in horses has been suggested previously but was not consistent in different mare populations (Nolan et al., 2017). Our present study compared the effects of administering daily extended photoperiod via blue LED light to control pregnancies in the same mares kept under identical conditions. Individual mare effects were thus excluded to a larger extent compared to studies using independent treatment and control groups. Furthermore, in the present study, blue LED light was applied from 8:00 throughout daylight time until 23:00. This ensured a constant and consistent light intensity for all mares with light masks, irrespective of changes in natural light intensity caused by the meteorological situation or an uneven light distribution in the stable. Our results support previous findings that suggest photoperiod influences gestation length, whether that is provided by the natural changes in seasonal daylength or artificial lighting programmes (Hintz et al., 1979; Hodge et al., 1982; Nolan et al., 2017). Equine gestation length is in part genetically determined and individual mares will repeatedly carry their foal for specific durations in multiple consecutive years (Kuhl et al., 2015) but the effects of blue LED light on gestation length are more pronounced than the genetic-associated differences among mares. Furthermore, in mares that conceive relatively late in the previous year’s breeding season, a longer than average gestation may result in delayed re-breeding. Whereas classical light programmes affect all mares in a group or stable, blue light-emitting diodes will allow targeting those mares individually.

The regulatory mechanisms through which increasing day length accelerates the onset of foaling remains open but the underlying pathways must be responsive to photoperiod. The initial stimulus for foaling is a shift from pregnenolone to cortisol synthesis in the fetal adrenal cortex. This shift is enabled by increasing responsiveness of the fetal adrenal gland to ACTH and indicates fetal maturation (Holtan et al. 1991; Silver and Fowden, 1994). Therefore, our results suggest stimulating effects of blue LED light on fetal maturation in horses. Enhanced maturation will then advance the fetal signals initiating the onset of foaling and result in earlier birth of the foal. Most likely, the natural seasonal increase in the duration of daylight will have a similar effect on fetal maturation and the timing of birth, resulting in a decrease in gestation length throughout the foaling season. This idea of a stimulatory effect of the blue LED light on fetal maturation is further supported by a higher neutrophil to lymphocyte (N/L) ratio in foals when mares had been exposed to blue LED light compared to foals from control pregnancies. Haematological changes are indicative of maturity of the foal and the N/L ratio is a useful index for assessing the foal’s maturity at birth. A threshold value of 2.5 indicates that the foal is fully mature (Jeffcott et al. 1982, Neuhauser et al., 2009). Changes in plasma progestagen concentration before and after foaling were identical to control pregnancies when mares were exposed to blue LED light. Peripartum changes in progestagen concentration were similar to previous studies (Haluska and Currie, 1988; Nagel et al., 2012), and indicate that endocrine preparation for foaling is accelerated but not negatively affected by blue LED light

About 90% of mares give birth during the night (Rossdale and Short, 1967; Heidler et al., 2004). Blue LED light did not influence the time that mares delivered their foals and thus did not shift foaling later into the night. However, the difference in hours of light that mares from blue LED light and control pregnancies were exposed to, either naturally or artificially via light masks, decreased as each study period progressed towards time of parturition. Thus, the effects of blue LED light on time of foaling during the night could not be fully evaluated in this study and may require comparison of treatments where more pronounced differences in photoperiod occur between blue LED light and control pregnancies at time of foaling.

Although our study cannot determine how the equine fetus is able to perceive changes in external light, the markedly shorter hair in foals born to light-treated mares clearly demonstrates that external light effects do reach the fetus. A shorter hair coat in foals born to mares wearing light masks has been demonstrated recently by Nolan et al. (2017) but in that study was not associated with a reduction in gestation length. Maternal melatonin is transferred to the fetus in sheep (Yellon and Longo, 1988), rats (Klein, 1972) and non-human primates (Reppert et al., 1979) and melatonin receptors are expressed in the fetus (Ebling et al., 1989; Drew et al., 1998). It thus can be assumed that blue LED light reduced the duration of the nocturnal phase of melatonin release in pregnant mares and such changes were transferred to the fetus. Furthermore, in sheep external light to a certain extent reaches the pregnant uterus. In late gestation, changes in intrauterine light radiation throughout the day are present reflecting the changes in external daylight. Thus, the sheep fetus is directly exposed to light-dark transitions at dawn and dusk (Parraguez et al., 1998). It is, however, unlikely that the blue LED light directed at one eye of the late pregnant mare is directly perceived by its fetus.

Foals born to mares exposed to blue LED light were slightly smaller than control foals but there were no differences in foal birth weights between groups. Whereas equine fetal weight gain increases to a maximum near term and is determined mainly by developing muscles and soft tissues, growth in length of the bones is maximal in months 8 and 9 of gestation (Platt, 1987, 1984). Therefore, the height differences between foals born from blue LED light-treated and control pregnancies are most likely caused by effects stimulated at the beginning or throughout the phase when mares were exposed to blue LED light and not caused by the shorter gestation. Birthweight has an ongoing influence on bodyweight in Thoroughbred race horses (Brown-Douglas et al. 2005). Foals with higher birthweights are heavier throughout life, and have a better racing performance compared to horses born with a low birthweight (Platt, 1987). A similar birthweight in foals from blue LED light and control pregnancies suggests no effects of treatment on the foals’ future performance as race horses.

In cattle, IgG content in colostrum changes with season and concentrations are higher in August and September than in other months of the year (Gay et al., 1983; Shearer et al., 1992). To the best of our knowledge, seasonal changes in colostral IgG content have not been studied in horses, but based on the findings in cattle, we hypothesized that an artificially extended photoperiod increases IgG transfer to equine colostrum. This hypothesis was not verified and, in contrast, IgG content in colostrum of mares exposed to blue LED light was slightly lower than in the same mares after control pregnancies Nevertheless, in both groups of mares, colostrum quality was in an adequate range to ensure sufficient IgG transfer to their newborn foals (Chavatte-Palmer et al., 2001) as indicated by semiquantitative IgG test.

In the mares, hair re-growth in an area shaved at the start of the experiment in December was less pronounced after exposure to blue LED light compared to control pregnancies while hair length in a non-shaved area did not differ between treatments. Effects of blue LED light on hair coat changes in horses depend on the time of the year when the light programme is started. When initiated one month before the winter solstice, blue LED light accelerated coat shedding in outdoor living horses but was without effect when initiated one month after the winter solstice (O’Brien et al., 2020). Exposure of mares to blue LED light in our studies began closer to the winter solstice than in the previous study (O’Brien et al., 2020). Although blue LED light therefore was without effect on overall hair length, it still reduced hair re-growth in mares to some extent. This difference is most likely associated with differences in prolactin secretion caused by the exposure of mares to blue LED light. On the days before foaling, prolactin concentration in plasma of mares tended to be elevated when mares were exposed to blue LED light compared to controls. Similar to our findings in mares exposed to blue LED light, either an artificial light programme providing a summer photoperiod or constant light increased plasma prolactin concentrations in pregnant ewes (Bassett, 1992; Parraguez et al., 1996). Effects of artificial light on maternal prolactin release in sheep were, however, more pronounced than in horse mares of the present study. In agreement with previous reports from our group (Heidler et al., 2003), in the present study prolactin concentration increased markedly at foaling. It thus cannot be excluded that the marked physiological prolactin increase associated with the onset of lactation in horses (Heidler et al., 2003) may in part have masked effects of blue LED light on prolactin release.

In conclusion, foaling can be advanced with head-worn masks emitting blue LED light to a single eye without negative effects on foal maturity. Blue LED light resulted in the birth of slightly smaller but more mature foals compared to controls. Furthermore, foals born to blue LED light-treated mares had a shorter hair coat than control foals, demonstrating that artificial light directed at the mare does reach its fetus and accelerates fetal maturation.

## Acknowledgements

The authors are grateful to Ronny Voigt and his team at the Neustadt (Dosse) *Hauptgestüt* for diligent care of mares and foals, to Julia Maderner for expert support with hormone analysis and to Christiane O’Brien for expert technical assistance with the Equilume light masks.

